# Enteric Pathogen Diversity in Infant Foods in Low-income Neighborhoods Of Kisumu, Kenya

**DOI:** 10.1101/493213

**Authors:** Kevin Tsai, Sheillah Simiyu, Jane Mumma, Rose Evalyne Aseyo, Oliver Cumming, Robert Dreibelbis, Kelly K. Baker

## Abstract

Pediatric diarrheal disease remains the 2nd most common cause of preventable illness and death among children under the age of five, especially in Low and Middle-Income Countries (LMICs). However, there is limited information regarding the role of food in pathogen transmission due to measuring infant food contaminations in LMICs. For this study, we examined the frequency of enteric pathogen occurrence and co-occurrence in 127 weaning infant foods in Kisumu, Kenya using a multi-pathogen rt-PCR diagnostic tool, and assessed household food hygiene risk factors for contamination. Bacterial, viral, and protozoa enteric pathogen DNA and RNA were detected in 62% of the infant weaning food samples collected, with 37% of foods containing more than one pathogen type. Multivariable generalized linear mixed model analysis indicated type of infant food best explained the presence and diversity of enteric pathogens in infant food, while most household food hygiene risk factors considered in this study were not significantly associated with pathogen contamination. Specifically, cow’s milk was significantly more likely to contain a pathogen (adjusted Risk Ratio=14.4; 95% Confidence Interval (CI) 1.78-116.1) and contained 2.35 more types of pathogens (adjusted Risk Ratio=2.35; 95% CI 1.67-3.29) than porridge. Our study demonstrates that infants in this low-income urban setting are frequently exposed to diarrhoeagenic pathogens in food and suggests that interventions are needed to prevent foodborne transmission of pathogens to infants.

**Importance:** Food is acknowledged as an important pathway for enteric pathogen infection in young children. Yet, information on enteric pathogen contamination in food in low-and-middle income settings is lacking, especially with respect to weaning foods given to young infants. This study assessed which food-related risk factors were associated with increased presence of and diversity in twenty-seven types of enteric pathogens in a variety of foods provided to infants between three and nine months of age in a low-income neighborhood of Kisumu, Kenya. Feeding infants cow milk emerged as the most important risk factor for food contamination by one or more enteric pathogens. The results indicate public health interventions should focus on improving cow milk safety to prevent foodborne pathogen transmission to infants. However, more research is needed to determine whether infant milk contamination was caused by caregiver hygiene practices versus food contamination passed from upstream sources.

## Introduction

Even though pediatric diarrheal diseases are declining worldwide, they remain the 2nd most common cause of preventable illness and death among children under the age of five [(1)]; responsible for approximately 800 million illnesses and 800,000 deaths in 2010. Approximately 90% of this disease burden is concentrated in children under the age of 5 in low- and middle-income countries (LMICs) [(2, 3)]. Diarrheal infections are caused by a diverse range of enteric pathogens that infect children as early as birth [(4)]. Children infected with enteric pathogens can potentially suffer long-term adverse effects to their physical and cognitive development and future socio-economic status [(5, 6)].

There is increasing recognition that consumption of pathogen-contaminated food is an important exposure pathway for diarrheal disease in children in LMICs [(7, 8)]. An estimated 582 million cases of illness, death, or disability adjusted life years are attributed to contaminated food each year globally, with young children and Africans bearing most of the foodborne disease burden [(2)]. The risk of diarrheal disease typically increases as infants transition from exclusive breastfeeding to consumption of weaning foods and water due to both decreases in passive protection from maternal breastmilk and more exposure to contaminated food [(9, 10)]. Many caregivers worldwide struggle to exclusively breastfeed up to six months of age, resulting in infants being provided weaning food instead of breast milk before 6 months of age [(11–13)]. Thus, premature transition from exclusive breast feeding to weaning foods may be especially important as one of the earliest causes of enteric infection [(14)]. Little is known about how often infants in LMICs are exposed to pathogens via food, and which risk factors should be targeted to reduce food-related exposure of children to enteric pathogens. In addition, infants’ diets become more diversified as they develop, and each of these additional food types may pose different exposure risks for different enteric pathogens [(15)]. More evidence is needed to understand which risk factors should be targeted to reduce food-related exposure of children to enteric pathogens in LMIC settings [(16)].

While foodborne transmission of enteric pathogens into the food supply chain is rigorously monitored in high-income countries (HICs) via regulatory authorities [(17)], food safety is frequently not monitored and regulated well in LMICs [(7)]. Many common infant weaning foods, like cow’s milk, are sourced from outside the household. Unsanitary and unregulated farm and market practices can result in contamination of milk by human or animal feces, well before entry to the household [(18)]. Reliance on unsanitary water to prepare weaning foods is common in LMICs [(10)]. In addition to sub-optimal water and market food supplies, insufficient hand washing and sterilization of food preparation areas, improper cooking temperature of infant food, storage of perishable foods at ambient temperatures, and storage of food in containers open to flies [(7, 19, 20)] can introduce additional microbial contamination in the household.

This study aimed (1) to describe the frequency and diversity in enteric pathogen contamination of infant weaning foods in low-income, neighborhoods of Kisumu, Kenya, and (2) to identity the leading environmental conditions and behaviors that contribute to pathogen presence and absence and higher pathogen diversity. The methodologies described and applied in this paper could be used in future research on foodborne illnesses in LMICs. Furthermore, our findings could be used to inform public health and healthcare professionals as a basis for prevention of pediatric diarrheal diseases in LMICs.

## Results

### Demographics of Caregivers/Infants and Household Hygiene Conditions

A total of 127 households (caregivers/infant dyads) participated in this study. Seventy-seven households were enrolled in January following the initial census and recruitment of all children between 3 and 9 months of age, and another 30 and 20 households were enrolled in March and May, respectively (Table 2). The study population was comprised of 45% male and 54% female infants born between March 2016 to December 2016. Most infants of caregivers were over 6 months old (76%). Among the caregivers who provided the study samples, 83% were married. Half of caregivers (50%) had only a primary education, whereas 21 *%* had some secondary education and 30% completed secondary education. Almost half (47%) of the caregivers who reported their employment status were unemployed (Table 2). There was variability in number of households enrolled across the four CHV catchment areas due to differences in number of eligible infants for recruitment, inability to locate the caregivers after census, refusals to participate, ability to verify infant’s age, or refusal to provide food samples after consent and participation in the survey.

**Table 1:**
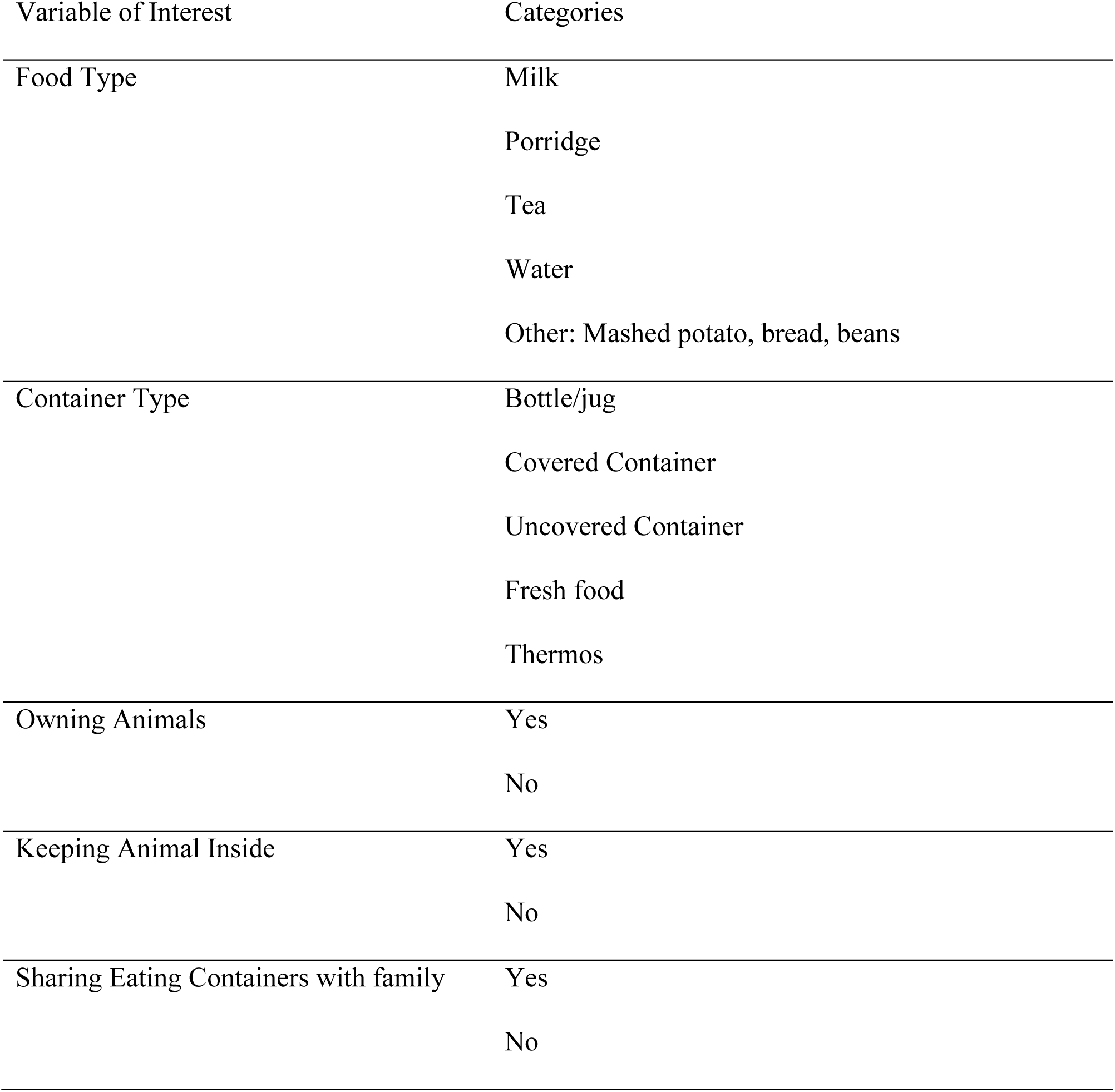
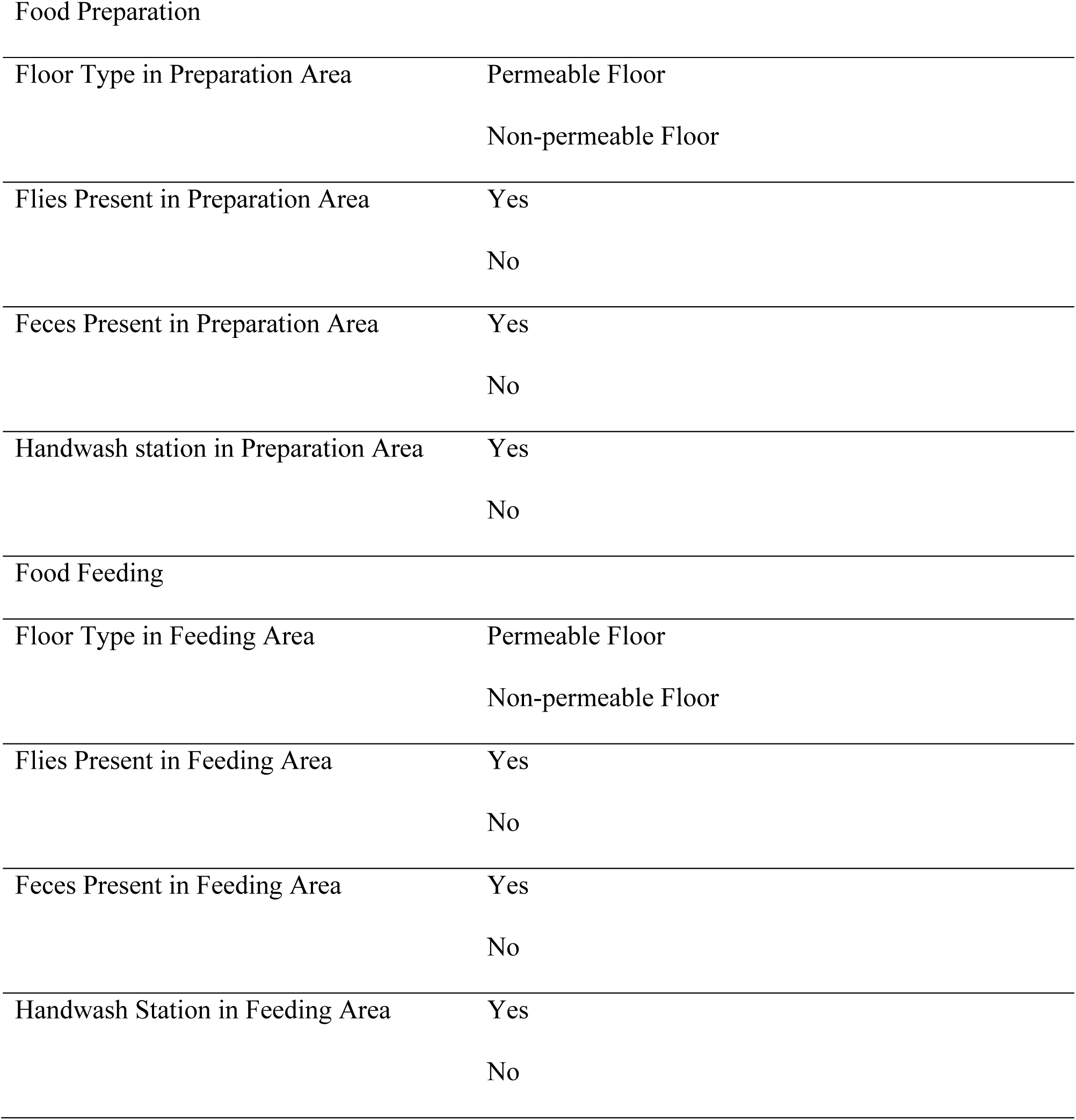
Variables representing Risk Factors for Food Contamination by Enteric Pathogens

| Variable of Interest | Categories |
| --- | --- |
| Food Type | Milk |
|  | Porridge |
|  | Tea |
|  | Water |
|  | Other: Mashed potato, bread, beans |
| Container Type | Bottle/jug |
|  | Covered Container |
|  | Uncovered Container |
|  | Fresh food |
|  | Thermos |
| Owning Animals | Yes |
|  | No |
| Keeping Animal Inside | Yes |
|  | No |
| Sharing Eating Containers with family | Yes |
|  | No |
| Food Preparation |  |
| Floor Type in Preparation Area | Permeable Floor |
|  | Non-permeable Floor |
| Flies Present in Preparation Area | Yes |
|  | No |
| Feces Present in Preparation Area | Yes |
|  | No |
| Handwash station in Preparation Area | Yes |
|  | No |
| Food Feeding |  |
| Floor Type in Feeding Area | Permeable Floor |
|  | Non-permeable Floor |
| Flies Present in Feeding Area | Yes |
|  | No |
| Feces Present in Feeding Area | Yes |
|  | No |
| Handwash Station in Feeding Area | Yes |
|  | No |

**Table 2.**
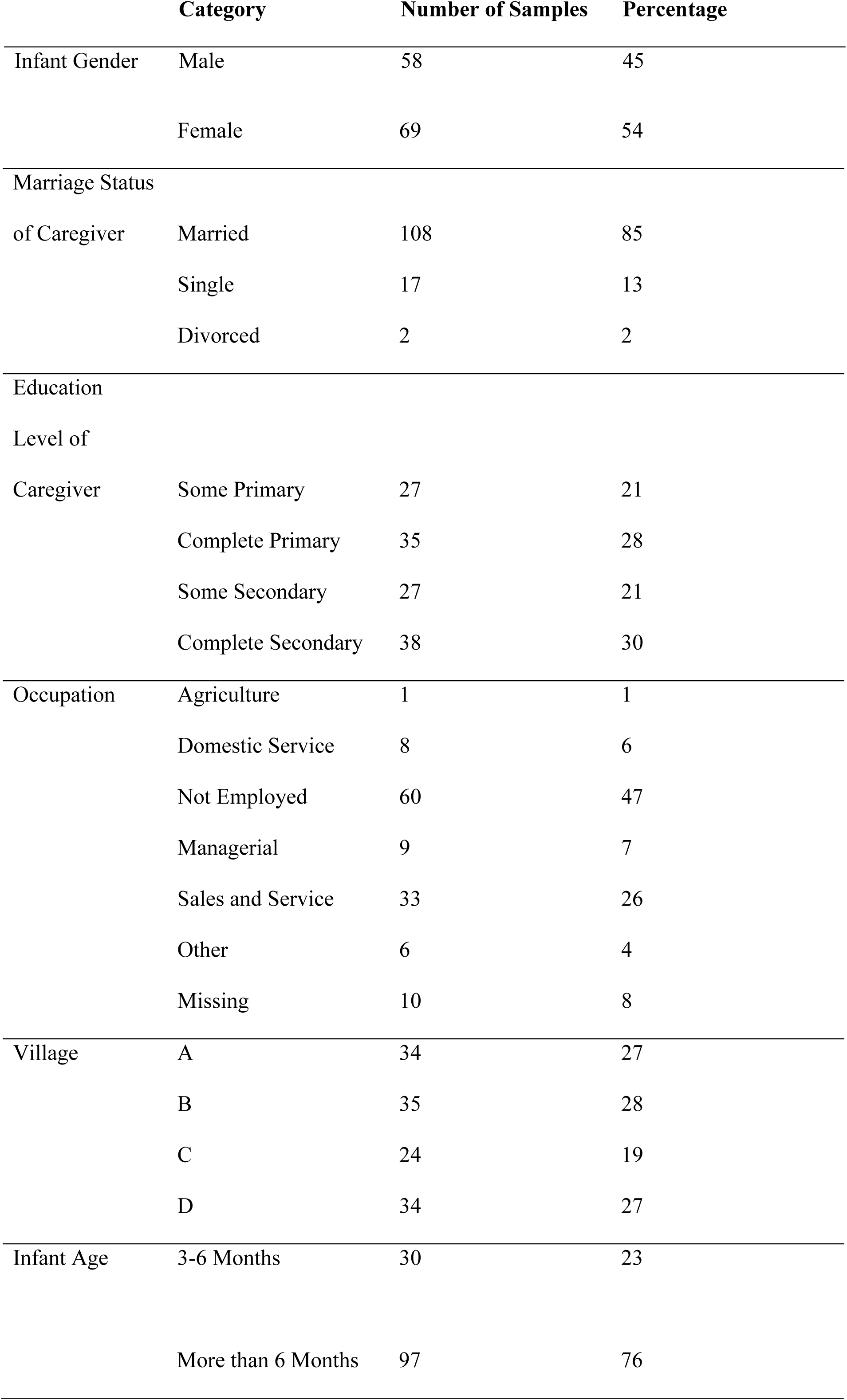
Socio-economic demographic statistics for 127 caregivers and infant dyads in Kisumu

The most common infant food types were porridge and cow’s milk, followed by tea, and “other” food (example: flour bread, mashed potatoes, or beans) (Table 3). Food type did not vary for infants < 6 months of age versus those older than 6 months (chi-square, p=0.12) when food types were categorized as milk, porridge, and non-milk/porridge (tea, water, flour bread, mashed potatoes, or beans were grouped due to low frequencies per category). Most of the households did not have a handwashing station in their food preparation and/or feeding area. Flies were observed in one third of household food preparation and feeding areas, and animal feces were observed in 8% of household food preparation and feeding areas. Non-permeable floors in food preparation and feeding areas were most common.

**Table 3:**
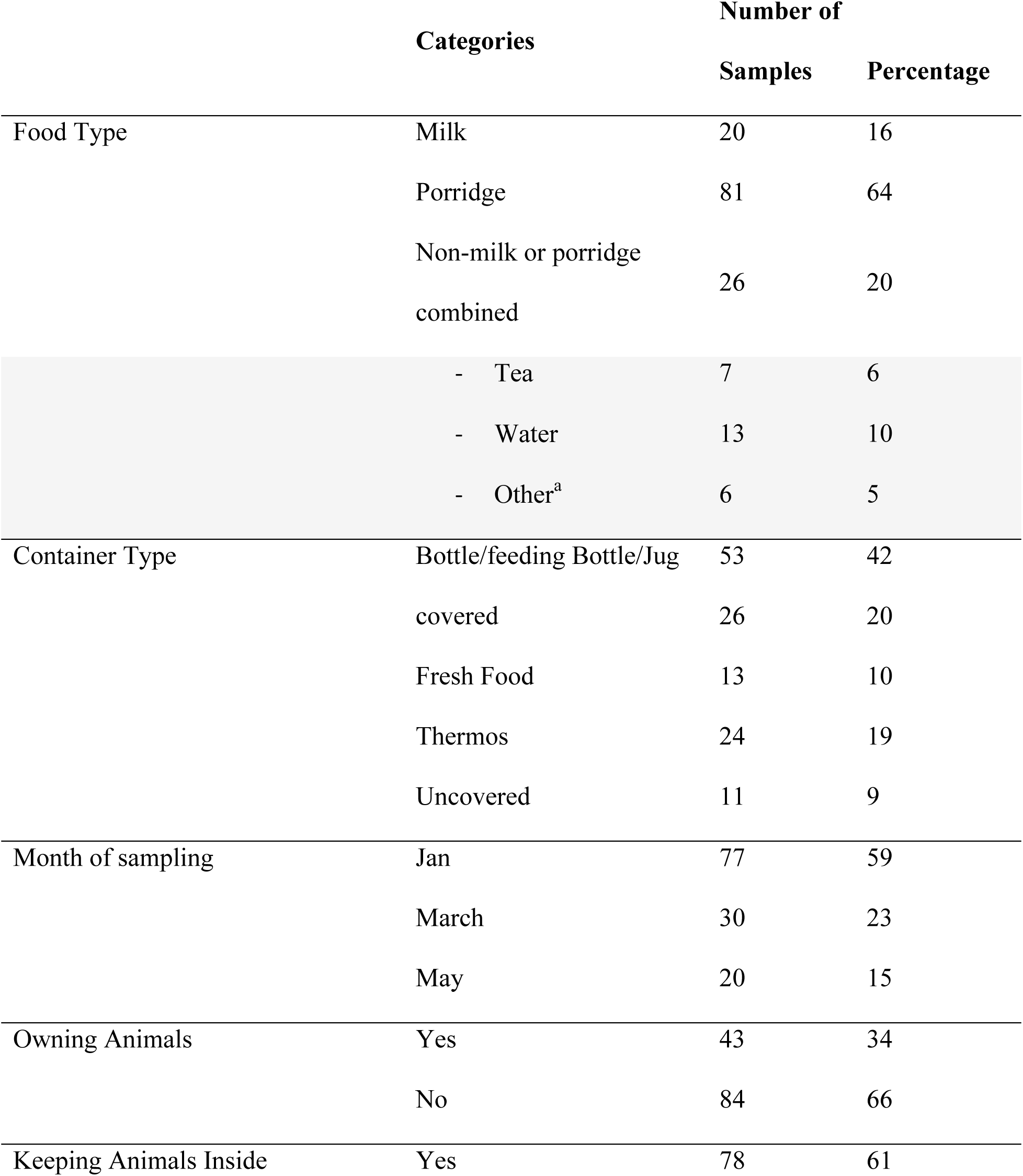
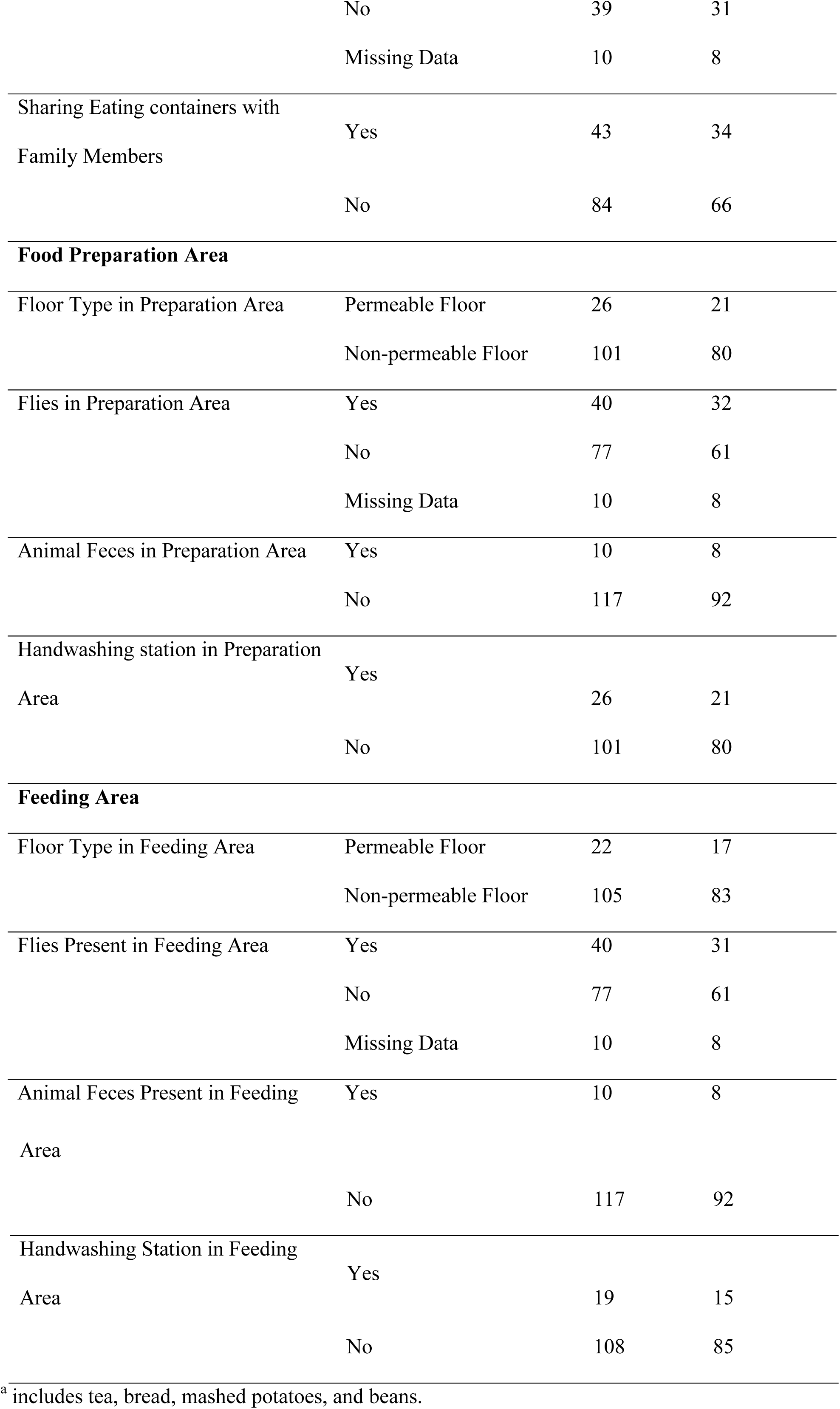
Food Contamination Risk Factor Statistics for households in Kisumu

### Pathogen Distribution and Diversity in Infant Weaning Foods

Assessment of the quality of DNA and RNA extracted from infant food is reported in Supplemental Table S2. DNA and RNA of 13 different types of bacterial, viral, and protozoa enteric pathogens was detected in 79 of the 127 (62%) infant weaning food samples collected over the three-month span (Table 4). The most commonly detected pathogens were *Aeromonas* (20%), *Enterohemorrhagic E. coli* 0157 (*EHEC*) (17%), *Enteropathogenic E. coli* (*EPEC*) (17%), *Enterotoxigenic E. coli* (*ETEC*) (13%), *Adenovirus 40/41* (12%), and *non-parvum/non-hominus Cryptosporidium spp*. (10%), with 8 other pathogens occurring in less than 10% of overall infant weaning food samples (Table 4). Infant food samples collected during March have higher raw contamination rate than food collected during January and May (90%, 52%, and 60%, respectively). ETEC and *non-parvum* or *hominus Cryptosporidium spp* were detected frequently in January. In March, detection frequencies for Adenovirus 40/41, EPEC, EHEC 0157, STEC, EIEC/*Shigella spp*., and *C. difficile* were highest. *Aeromonas* were detected frequently in May. A median of 1 pathogen per sample (standard deviation of 1.58; range of 0 to 9 pathogen) was detected, with 37% of foods being co-contaminated by more than 1 pathogen type.

**Table 4:** Pathogen Presence by Month.

|  | Overall<br>(Total=127) | January (Total=77) | March (Total=30) | May (n=20) |
| --- | --- | --- | --- | --- |
| Number (Percentage)<br>positive | 79 (62) | 40 (52) | 27 (90) | 12 (60) |
| Virus |  |  |  |  |
| Adenovirus 40/41 | 15 (12) | 3 (3) | 10 (33) | 2 (10) |
| Adenovirus Hexon | 6 (5) | 1 (1) | 3 (10) | 2 (10) |
| Norovirus | 9 (7) | 4 (5) | 3 (10) | 2 (10) |
| Sapovirus | 1 (1) | 1 (1) | 0 (0) | 0 (0.0) |
| Bacteria |  |  |  |  |
| EAEC | 6 (5) | 4 (5) | 2 (7) | 0 (0.0) |
| EPEC | 21 (17) | 3 (4) | 15 (50) | 3 (15) |
| ETEC | 17 (13) | 13 (17) | 3 (10) | 1 (5) |
| EHEC 0157 | 21 (17) | 0 (0) | 21 (70) | 0 (0.0) |
| STEC | 5 (4) | 0 (0) | 5 (17) | 0 (0.0) |
| EIEC/Shigella | 7 (6) | 4 (5) | 3 (10) | 0 (0.0) |
| Aeromonas | 25 (20) | 12 (16) | 5 (17) | 8 (40) |
| *B. Fragilis* | 1 (1) | 0 (0) | 0 (0) | 1 (5) |
| *C. difficile* | 11 (9) | 5 (7) | 5 (17) | 1 (5) |
| Protozoa |  |  |  |  |
| *Cryptosporidium spp.* | 13 (10) | 10 (13) | 2 (7) | 1 (5) |

<sup>a</sup>. No detection for Astrovirus, Rotavirus, *Salmonella\_enterica*, *H. pylori*, *Vibrio Cholerae*, *Vibrio parahaemolytic*, *Giardia lamblia*, *Cryptosporidium hominus*, *Cryptosporidium parvum*, *E. histolytica*, *A. Lumbricoides*, *N. americanus*, *S. Sterocoralis*, *T. trichiura*

### Risk Factors for Enteric Pathogens presence in Infant Weaning Food

Food type, the infant sharing eating containers with other family members, and feces in preparation area were associated at p<0.3 with presence of any pathogen in the bivariate analysis, and were included in the multivariable analysis (Table 5). Sharing an eating container did not improve model fit and was removed. In the final multivariable model, cow milk was significantly more likely to contain an enteric pathogen when compared with porridge, but non-milk/porridge foods were not statistically different from porridge. Pathogens were detected twice as often in milk (95%, n=19/20) as porridge (56%, n=45/81) and non-milk/porridge foods (56%, n=15/26). Observation of feces in preparation area was statistically associated with a lower risk of pathogen presence compared to feces not being observed.

**Table 5:**
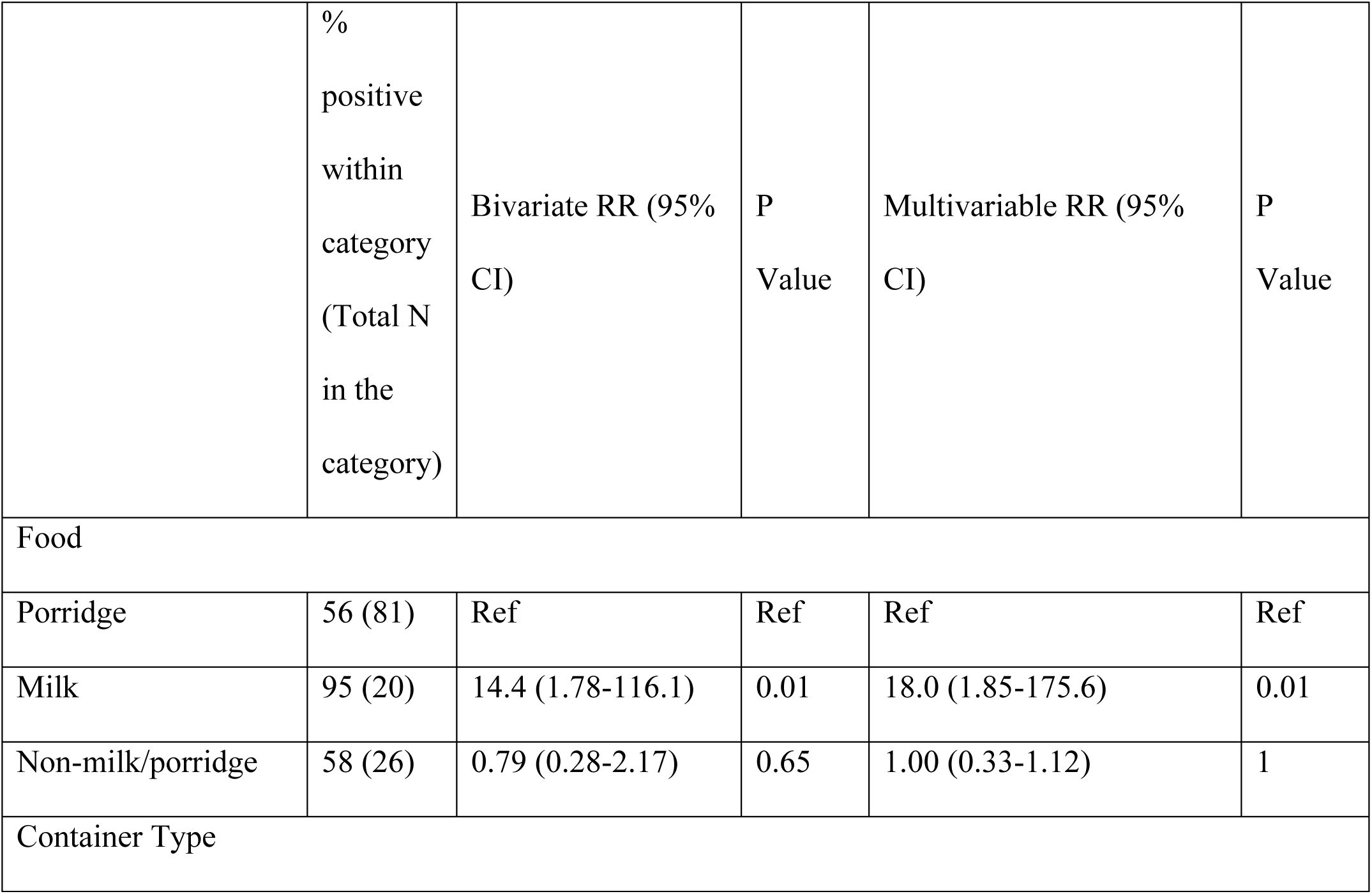
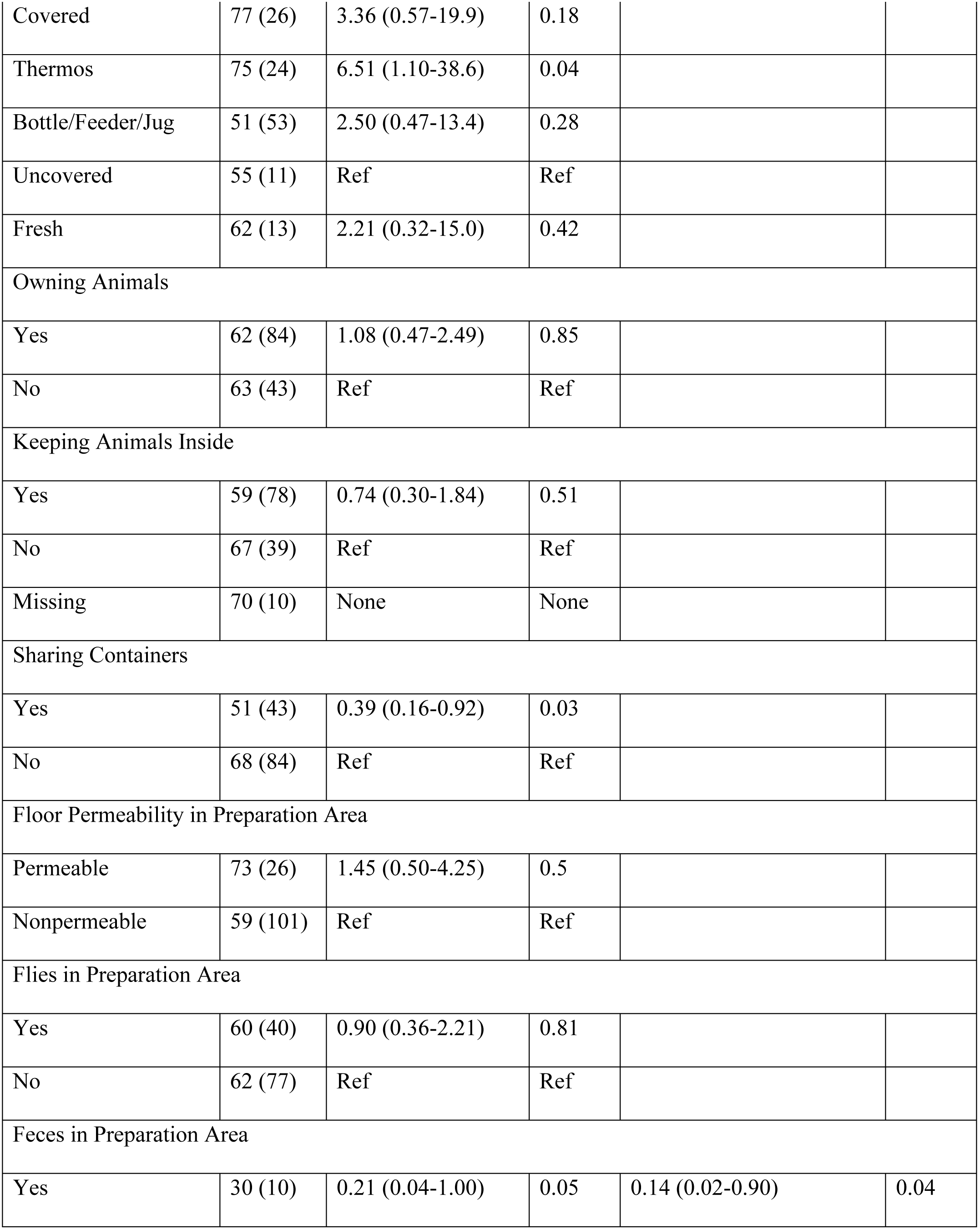
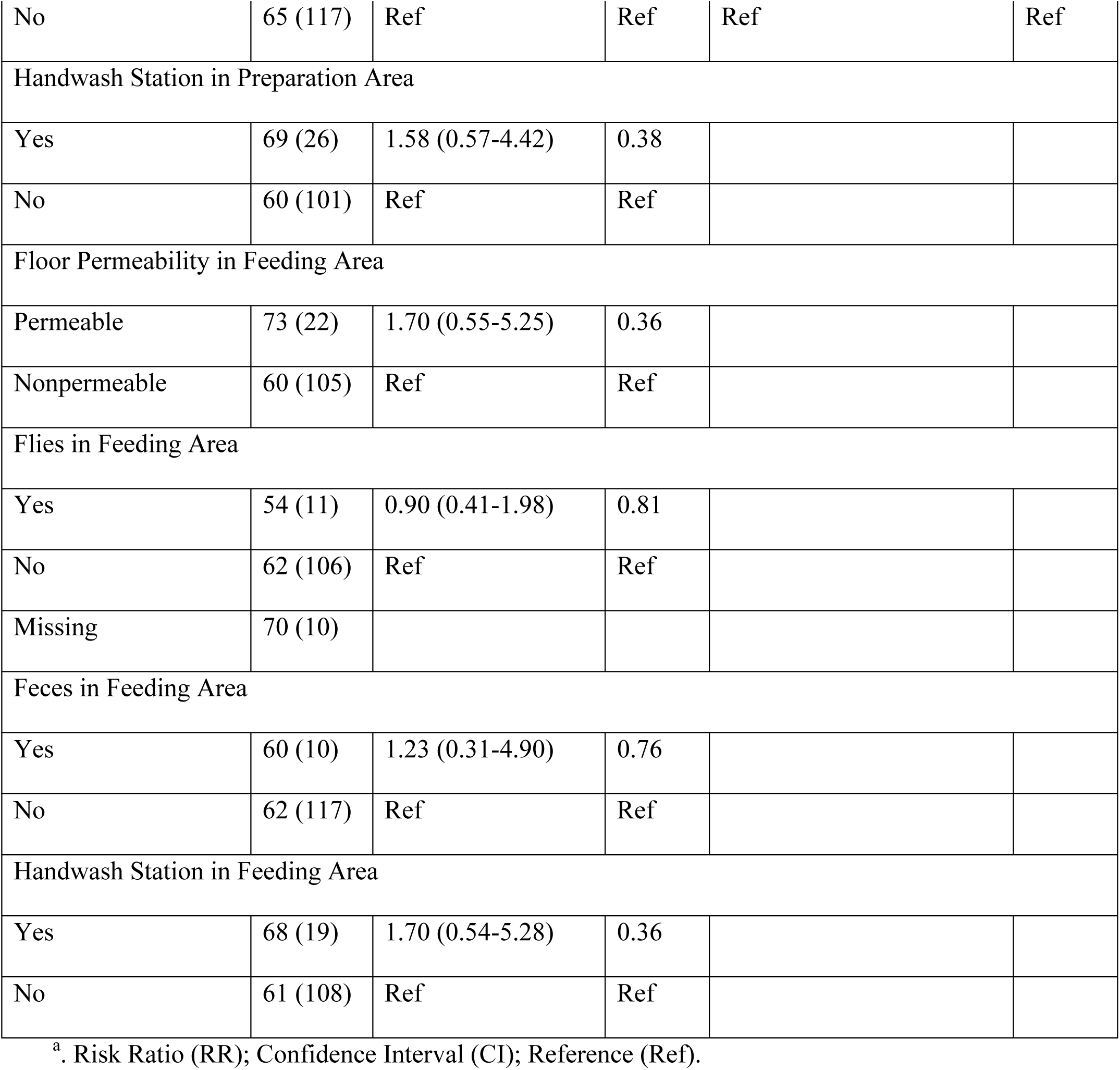
Bivariate and multivariable generalized linear mixed models of food contamination risk factors and enteric pathogen presence in infant weaning foods

|  | %<br>positive<br>within<br>category<br>(Total N<br>in the<br>category) | Bivariate RR (95%<br>CI) | P<br>Value | Multivariable RR (95%<br>CI) | P<br>Value |
| --- | --- | --- | --- | --- | --- |
| Food |  |  |  |  |  |
| Porridge | 56 (81) | Ref | Ref | Ref | Ref |
| Milk | 95 (20) | 14.4 (1.78-116.1) | 0.01 | 18.0 (1.85-175.6) | 0.01 |
| Non-milk/porridge | 58 (26) | 0.79 (0.28-2.17) | 0.65 | 1.00 (0.33-1.12) | 1 |
| Container Type |  |  |  |  |  |
| Covered | 77 (26) | 3.36 (0.57-19.9) | 0.18 |  |  |
| Thermos | 75 (24) | 6.51 (1.10-38.6) | 0.04 |  |  |
| Bottle/Feeder/Jug | 51 (53) | 2.50 (0.47-13.4) | 0.28 |  |  |
| Uncovered | 55 (11) | Ref | Ref |  |  |
| Fresh | 62 (13) | 2.21 (0.32-15.0) | 0.42 |  |  |
| Owning Animals |  |  |  |  |  |
| Yes | 62 (84) | 1.08 (0.47-2.49) | 0.85 |  |  |
| No | 63 (43) | Ref | Ref |  |  |
| Keeping Animals Inside |  |  |  |  |  |
| Yes | 59 (78) | 0.74 (0.30-1.84) | 0.51 |  |  |
| No | 67 (39) | Ref | Ref |  |  |
| Missing | 70 (10) | None | None |  |  |
| Sharing Containers |  |  |  |  |  |
| Yes | 51 (43) | 0.39 (0.16-0.92) | 0.03 |  |  |
| No | 68 (84) | Ref | Ref |  |  |
| Floor Permeability in Preparation Area |  |  |  |  |  |
| Permeable | 73 (26) | 1.45 (0.50-4.25) | 0.5 |  |  |
| Nonpermeable | 59 (101) | Ref | Ref |  |  |
| Flies in Preparation Area |  |  |  |  |  |
| Yes | 60 (40) | 0.90 (0.36-2.21) | 0.81 |  |  |
| No | 62 (77) | Ref | Ref |  |  |
| Feces in Preparation Area |  |  |  |  |  |
| Yes | 30 (10) | 0.21 (0.04-1.00) | 0.05 | 0.14 (0.02-0.90) | 0.04 |
| No | 65 (117) | Ref | Ref | Ref | Ref |
| Handwash Station in Preparation Area |  |  |  |  |  |
| Yes | 69 (26) | 1.58 (0.57-4.42) | 0.38 |  |  |
| No | 60 (101) | Ref | Ref |  |  |
| Floor Permeability in Feeding Area |  |  |  |  |  |
| Permeable | 73 (22) | 1.70 (0.55-5.25) | 0.36 |  |  |
| Nonpermeable | 60 (105) | Ref | Ref |  |  |
| Flies in Feeding Area |  |  |  |  |  |
| Yes | 54 (11) | 0.90 (0.41-1.98) | 0.81 |  |  |
| No | 62 (106) | Ref | Ref |  |  |
| Missing | 70 (10) |  |  |  |  |
| Feces in Feeding Area |  |  |  |  |  |
| Yes | 60 (10) | 1.23 (0.31-4.90) | 0.76 |  |  |
| No | 62 (117) | Ref | Ref |  |  |
| Handwash Station in Feeding Area |  |  |  |  |  |
| Yes | 68 (19) | 1.70 (0.54-5.28) | 0.36 |  |  |
| No | 61 (108) | Ref | Ref |  |  |

<sup>a</sup>. Risk Ratio (RR); Confidence Interval (CI); Reference (Ref).

Food type, handwashing station in preparation area, sharing eating containers with family members, and owning animals were associated with higher pathogen diversity at p<0.3 in bivariate analysis and were considered in the multivariable model (Table 6). Food type was the only variable retained in the final model for explaining pathogen diversity. Pathogen diversity was 2.35 times higher in milk than in porridge, whereas non-milk/porridge foods trended towards lower levels of diversity.

**Table 6:**
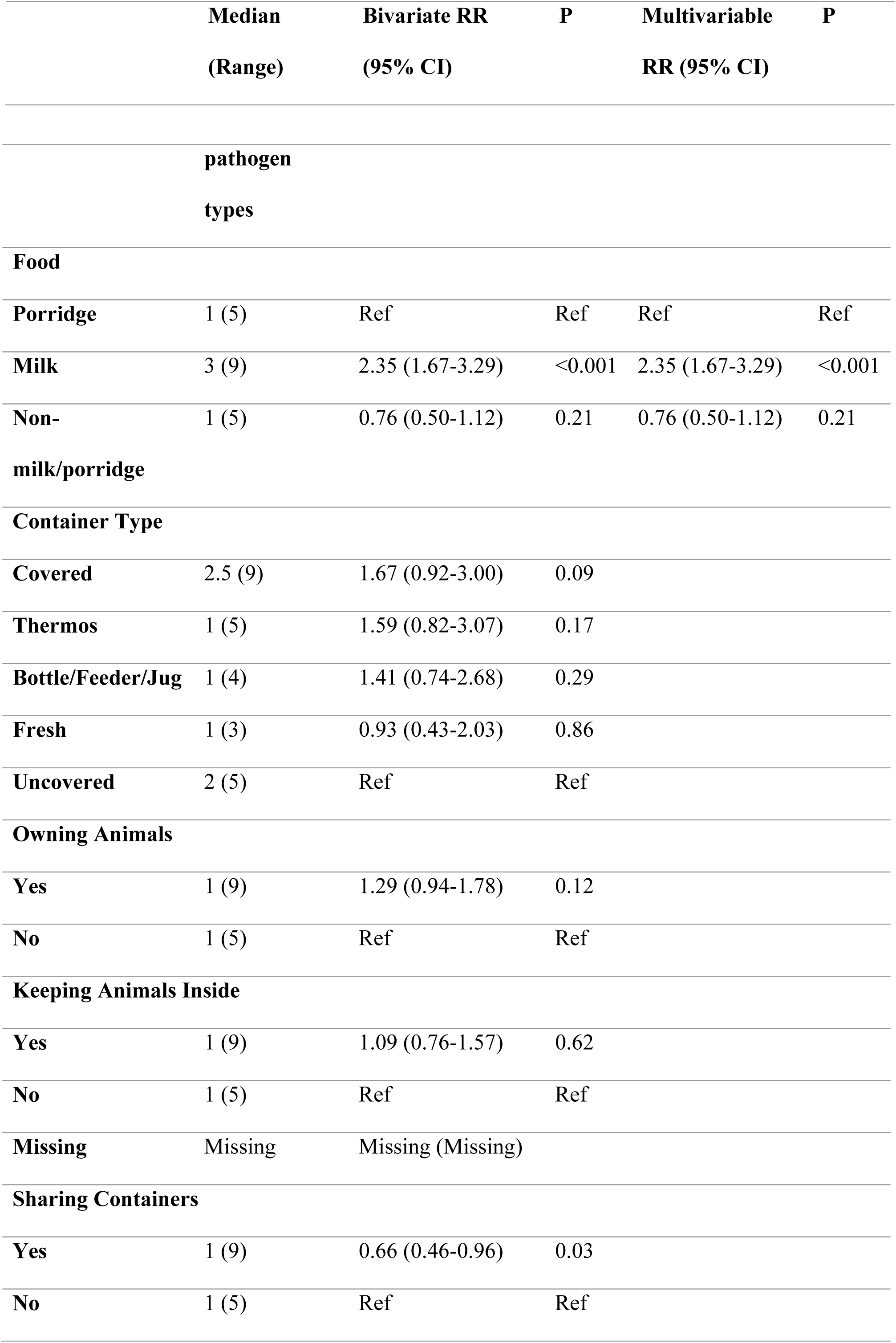
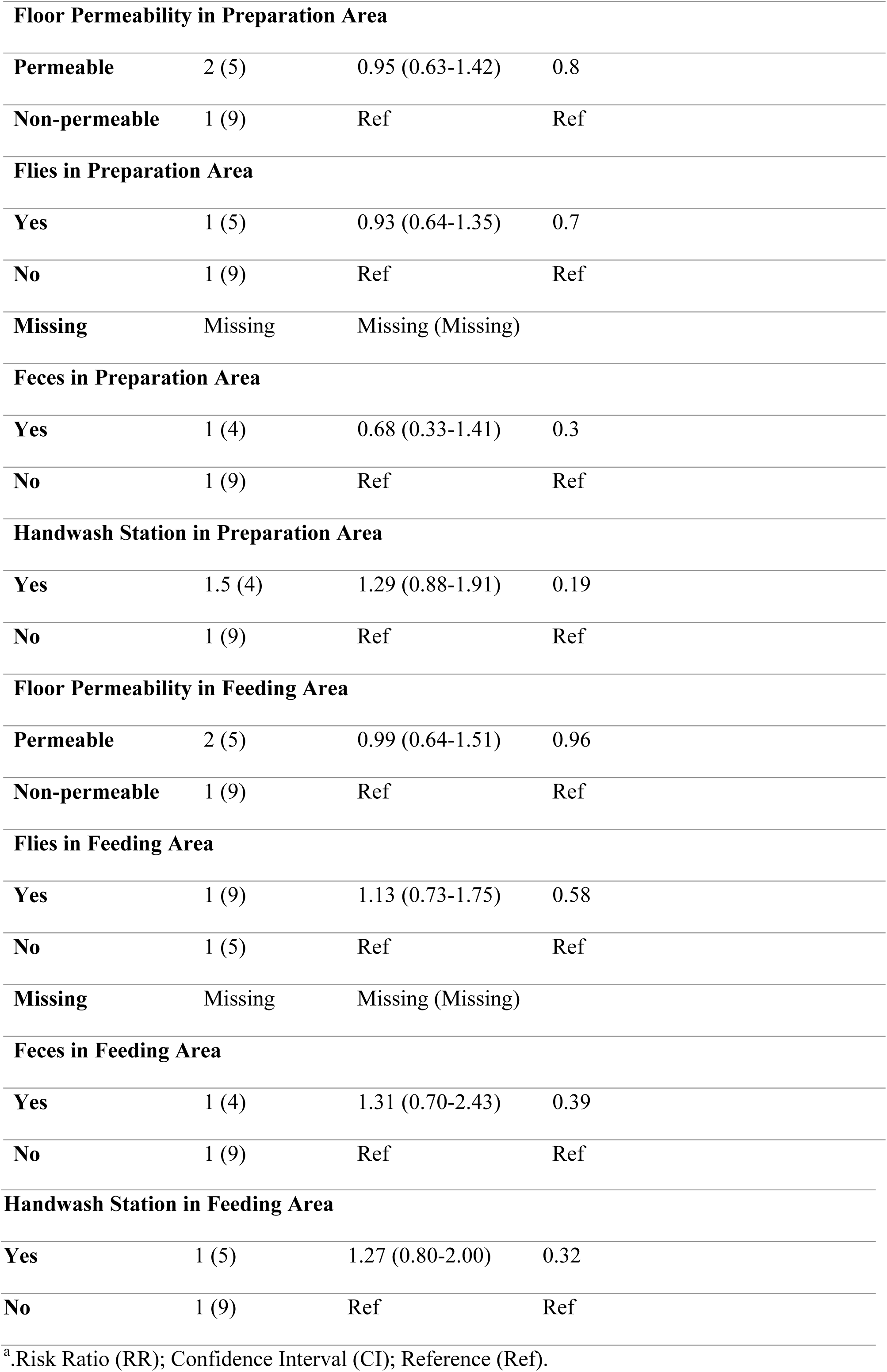
Bivariate and multivariable generalized linear mixed models of food contamination risk factors and enteric pathogen diversity in infant weaning foods.

## Discussion

Estimates of the importance of food as an enteric infection pathway for young children in LMICs are limited by the absence of primary data on food outbreaks and frequency of food contamination by enteric pathogens, especially with respect to weaning foods provided to infants [(17)]. This study demonstrated that infants as young as three months of age in informal settlements of Kisumu ingest food contaminated by a variety of different types of enteric pathogens. Our qPCR-based enteric pathogen detection frequency of 62% is similar to what has been reported for frequency of fecal indicator bacteria in infant food in similar high disease burden settings, such as Bangladesh (40% - 58%), Indonesia (45%), South Africa (70%), India (56%), and Peru (48%) [(2, 23–27)]. We expand upon these studies to show that a substantial number of infants ingest food contaminated by multiple types of enteric pathogens. Studies vary in their conclusions as to which pathogens cause the most foodborne enteric disease in LMICs, e.g. Norovirus, *Campylobacter spp., S. enterica, ETEC, EPEC, Giardia lamblia*, and *Shigella spp*. are all attributed with a substantial amount of foodborne illness or death [(2, 28)]. The etiology of foodborne disease may vary year to year, or month to month as suggested by our study, which has implications for ranking the priority foodborne pathogens in settings where outbreak or food monitoring information is limited. Many types of pathogens were detected in food during our 5-month study in Kisumu, with *Aeromonas* being the most common pathogen, followed by *EHEC O157, ETEC and EPEC*, and *human adenovirus 40/41. Aeromonas* is extremely common in the environment, including foods, but is not considered a priority foodborne pathogen [(29)]. However, *EHEC O157* is notorious as a deadly cause of foodborne epidemics, and the emergence of so many O157-positive food samples in March alone suggests there may have been a foodborne outbreak. March is the onset of the rainy season in Kisumu. The increased detection of multiple during this month may reflect an influence of seasonality on foodborne transmission risks in Kenya. This foodborne danger would have been missed had we sampled in a narrower timeframe.

We demonstrated that the risk of pathogen exposure for an infant can vary by type of weaning food, which has important implications for designing interventions. Cow’s milk was significantly more likely to be contaminated by one or multiple types of enteric pathogens compared to other common infant foods, such as porridge. For many urban and rural Kenya, raw milk is more affordable and accessible than pasteurized milk [(30)]. However, raw milk can be easily contaminated during production at farms by animal urine and feces, dirt, flies, adulteration with untreated water, and improperly cleaned containers [(31)]. In addition, the packaging, storage, distribution, and marketing of milk are not rigorously regulated and monitored in Kenya, leading to additional points where unhygienic conditions can introduce contamination [(32)]. Urban populations often encounter milk adulterated by water [(33, 34)]. Therefore, pasteurization of milk at the point of sale or during food preparation in the household may be critical for rendering milk safe to drink.

After sale, household food preparation, feeding, and storage conditions can contribute to new sources of infant food contamination [(19)]. In Kenya, milk is often consumed in liquid form, as well as is added to a variety of infant foods. Depending upon how the milk is provided to the infant, it may or may not receive proper treatment to eliminate microbial pathogens. If caregivers perceive milk to be safe due to prior pasteurization, they may not treat it further. If milk is added to infant foods, it may be reheated as a part of the cooking process or can be added to food after the preparation process without reheating. Milk is an optimal growth medium for bacteria and may be particularly sensitive to cross-contamination from unclean surface, hands, and flies or uncovered and unclean containers. Public health interventions targeting safety of milk products may be particularly effective for reducing foodborne diarrheal diseases in infants in LMICs.

Household water, sanitation, and hygiene (WASH) interventions have been suggested as keys in combating enteric pathogen transmission and infections [(16, 35, 36)]. While we found some domestic food hygiene risk factors were associated with enteric pathogen presence and diversity in bivariate analysis, most of these associations reduced in magnitude and did not improve model fit after adjustment for food type. Counterintuitively, observation of feces in the food preparation area – a rare situation to begin with - was associated with a lower likelihood of pathogen presence after model selection, rather than higher child food contamination as expected [(19)]. This association may be caused by unmeasured confounding factors or reactivity of some caregivers who were aware of the purpose of our visit. Timing household visits to coincide with food availability is logistically challenging unless caregivers store food for infants for prolonged periods of time. Thus, if food was not present during our first visit, we had to work carefully with households to time our follow-up visits to coincide with when they would have food for the child. Some caregivers may have reacted to the presence of feces in their preparation area before our visit and contaminated the infant’s food in the process of removing it. The inability to determine causality is a limitation of our cross-sectional design. The lack of association between other well-known risk factors of bacterial contamination in infant food could be caused by lack of statistical power to detect smaller effect sizes, although does not detract from the dominant role of food type in explaining pathogen detection. Analysis with a larger sample size are underway to improve knowledge about foodborne pathogen transmission in Kisumu.

One of the strengths of our study is that we examined food for pathogens, rather than bacterial indicators, using rigorous microbiological protocols to ensure quality of data was preserved from field labs in Kisumu to molecular labs in Iowa. Fecal indicators, which are typically used as a proxy for determining risk from fecal pathogens, are nonspecific and often do not correlate well with viral, bacterial, and protozoan pathogens [(37)]. Addressing the need for information on infant food contamination in LMICs required finding an effective microbial testing method that enabled quantitative and target-specific measuring of a broad array of the most common types of diarrhea pathogens in infant food. Even though qPCR is frequently applied for the quantitative detection of pathogen presence in foodborne outbreak analysis [(38–40)], it has not been widely applied in food samples in LMICs. Our methods are novel in their ability to detect a wide array of pathogens simultaneously. The qPCR approach is also a limitation, since qPCR-determined concentrations may detect non-infectious organisms that cannot cause disease. We are not certain what fraction of the PCR-detected pathogens are viable, viable-but-non-recoverable, or dead microbial organisms. However, the distinct variability in contamination patterns in infant food and the consistency with cow’s milk came up as a risk factor for pathogen presence suggests qPCR was a valid approach for identifying infant food risk factors.

Several challenges had to be overcome for measuring and analyzing infant weaning food contamination in this study, challenges which apply to many LMIC settings. First, multi-target enteric pathogen detection capability is limited in LMICs due to limited laboratory facilities, requiring samples to be sent to specialized labs for precise analysis. We minimized sample degradation risks by using a ZymoBIOMICS™ DNA/RNA extraction kit that allowed us to preserve samples at ambient temperature for storage, transport, and extraction [(41)]. This makes the method more desirable for use in laboratory-limited LMICs or any field-based scenario, as samples can be shipped to an equipped laboratory for processing with ease. The high rates of recovery of MS2 virus spiked in to samples before storage and transport confirmed that we experienced no loss in nucleic acid using this process. Second, the wide variety of physical and chemical properties of different food types makes optimization of microbial food testing protocols complex, especially if the goal is to measure multiple types of pathogens [(42)]. In addition, the presence of inhibitors can impact qPCR performance [(43)]. We pre-piloted all protocols to confirm that protocols for DNA and RNA recovery of spiked pathogens was not affected by food type, then rigorously evaluated each sample for signs of inhibition prior to qPCR analysis. Low inhibition rates and low variability in MS2 Ct values across all food types showed that Zymo extraction kit can produce high-quality nucleic acids free of inhibition from foods [(44–46)]. Third, pathogens may be present in food at concentrations that are lower than the methodological limit of detection, which results in misclassification of some pathogen-positive samples as uncontaminated. Pre-amplification increased the concentration of starting content before conducting the quantitative measurement step of PCR.

## Conclusion

Foodborne disease transmission of enteric pathogens may contribute substantially to the global diarrheal disease burden, yet receives limited attention. Our evidence highlights a need for more interventions targeting safe preparation and storage of infant foods, particularly high-risk foods such as milk. The ongoing Safe Start study in Kisumu is evaluating whether behavior improvements in caregiver food preparation, feeding, and storage behaviors can reduce enterococcus contamination in infant food and enteric pathogen infections in infants during weaning. Alongside interventions aiming to improve food hygiene practices of caregivers, interventions targeting hygienic milk handling and storage at the point of sale and among manufacturers may be needed to address upstream risks. The intersecting Market to Mouth study will contribute more information about the role of locally sold milk sources on pathogen contamination of infant food and the ability of the Safe Start intervention to mitigate enteric pathogen contamination passed via the food system.

## Materials and Methods

### Study Setting/Ethical Consideration

This exposure assessment study was conducted as part of formative research aimed at developing and testing an infant hygiene intervention to inform the development and evaluation of an infant weaning food hygiene intervention in Kisumu, Kenya. Kisumu is a city in the western region of Kenya, with a projected population of approximately 490,000 people by 2017 (Kisumu county integrated development plan 2013–2017). The study site includes four villages of a low-income peri-urban neighborhood in Kisumu. This infant weaning food hygiene intervention will be evaluated as part of the Safe Start study, a cluster-randomized controlled trial (Clinical Trials identifier: NCT03468114) involving Great Lakes University of Kisumu, Kenya (GLUK), the London School of Hygiene & Tropical Medicine (LSHTM), and the University of Iowa (UI). The study was approved by the scientific and ethical review committees at the GLUK (Ref. No. GREC/010/248/2016), LSHTM (Ref. No. 14695), and UI (IRB ID 201804204).

### Study Design

A total of eight community health volunteers (CHVs) who served the four neighborhoods in our study area facilitated the recruitment of participants. First, CHVs conducted a household census with the research team in December 2016 to generate a list of all infants less than nine months of age that were living in each CHV’s catchment area. Then, the list of households was randomly sorted, and in January CHV’s and enumerators approached each house to verify infant eligibility, obtain consent to participate in the study, and perform data collection, and food sampling. CHVs maintained surveillance of their respective catchment area through May 2017 to identify new infants as they became age-eligible, and to approach their caregivers about participation in the study.

Eligibility of a household was defined as having an infant between three and nine months of age, verified by reviewing the infant’s birth identity card, who was being fed supplemental food in addition to or in replacement of breastfeeding. Exclusion criteria included refusal to participate, inability to produce infant health card for verification of age, or caregiver reporting that the infant is exclusively breastfed and does not eat other food or liquid. Upon verification of eligibility and availability of food for sampling, consent to participate in the study was obtained from the child’s primary caregiver in the presence of the CHVs. The study was described in the caregiver’s natural language, and a signed copy of the consent form was left for her records.

### Data and Sample Collection

After agreeing to participate in the study, caregivers were interviewed about household status, their level of education; access to water, sanitation, and hygiene resources; and key infant weaning food preparation, storage, and feeding practices. Caregivers were then asked to provide approximately five grams of already-prepared infant food of any type fed to the child that day. The timing of food preparation for infants varied throughout the day, so the field team scheduled follow-up visits with households at times when food would be available. Food was placed into a sterile, labeled WhirlPak bag (Sigma-Aldrich, St. Louis, MO) by the caregivers, using whatever means (fingers, utensil) that the caregiver normally used for handling the child’s food. Food was placed on ice packs in a cooler and was transported to the laboratory for processing within six hours of collection.

### Nucleic Acid Extraction

All food samples were processed by following the manufacturer's instructions for the ZymoBIOMICS™ DNA/RNA extraction mini-kit (Zymo Research, Irvine, CA) for DNA & RNA parallel purification. A 250 mg food sample was measured into the Zymo-Shield tube, vortexed until blended, and stored at 4°C. Samples were then transported in a cooler at ambient temperature to the University of Iowa for the remainder of the extraction. A subset of samples (n=77) were spiked with 5 μL of 1.8*10^9^ CFU/mL of live bacteriophage MS2 to serve as an extrinsic process control to assess for RNA degradation as a function of storage and transport conditions. Once purified DNA/RNA was obtained, it was stored at −80°C until further analysis.

### Inhibition Screening/Preamplification

DNA and RNA extracts from the samples (6 μL each) were screened for evidence of inhibition with the QuantiFast Pathogen PCR +IC Kit and QuantiFast Pathogen qRT-PCR+ IC kit (Qiagen, Hilden, Germany) on a QuantStudio real-time PCR system (Thermo Fisher, Waltham, MA). A total of 77 samples were screened for inhibition. Inhibition was defined as having amplification of the RNA internal control over cycle threshold value (CT value) of 34 in a sample according to the manufacturer’s protocol. QuantiFast Pathogen qRT-PCR +IC Kit (Qiagen, Hilden, Germany) was used for pre-amplification PCR. For each sample, a total volume of 12 μL of DNA and RNA extract (6 μL each) was mixed with a master-mix containing 5 μL of 5x Quantifast Pathogen MM, 0.25 μL of 100x Quantifast Pathogen RT Mix, 0.5 μL of 50x high ROX dye solution, 0.15 μL of ultrapure 50 mg/mL BSA, 2.5 μL of 0.2x custom TaqMan pre-amplification primer and probe pool (Appendix 1), 2.5 μL of internal control assay, and 2.0 μL of internal control RNA. If extracts were determined to be inhibited during the inhibition screening, the inhibited extracts would undergo 1:10 dilution before mixing with the pre-amplification master-mix. The cycling conditions for the pre-amplification PCR were: holding stage of 50°C for 20 minutes and 95°C for 5 minutes, followed by 44 cycles of 95°C for 15 seconds and 60°C for 30 seconds [(21)]. Preamplification PCR was completed through an Eppendorf Thermocycler (Hamburg, Germany). All the completed pre-amplified samples would undergo a 1:10 dilution with nucleic acid-free water before proceeding to TaqMan quantitative PCR card analysis.

### TaqMan Array Card Analysis

Primers and probes for a total number of 37 gene targets of pathogen of interest in the TaqMan assays are listed in Supplemental Table S1. The Ag-Path-ID One-Step RT-PCR kit (Thermo Fisher, Waltham, MA) was used for the TaqMan card analysis. For each sample, 40 μL of re-amplified DNA/RNA extract (in 1 to 10 dilutions with nucleic acid-free water) was mixed with 50 μL of 2X RT-buffer, 4 μL of 25X AgPath enzyme, and 6 μL of nucleic acid-free water. All the TaqMan runs were completed in a ViiA7 instrument (Thermo Fisher, Waltham, MA), and the cycling conditions were: 45°C for 20 minutes and 95°C for 10 minutes, followed by 45 cycles of 95°C for 15 seconds and 60°C for 1 minute. Amplification of a pathogen-specific gene target was used to define a sample as positive for the presence of that pathogen; if multiple gene targets were used to detect different one type of pathogen (norovirus, *EAEC, EPEC, ETEC, STEC*) amplification of either gene resulted in a sample being considered positive. Two virulence gene indicators were used to detect pathogenic bacteria on the card, so in this manuscript samples were considered positive for the overall species of bacterial pathogen if either gene was detected.

### Data Analysis

There were two primary outcomes assessed during analysis. First, a binary indicator was defined based on the presence of one or more target pathogens detected in the sample (*any-path*). Second, pathogen diversity was calculated by summing the total number of target pathogens types detected in the sample (*sum-path*). Caregiver education level and sampling month were selected *a priori* as potential confounders of infant health and caregiver food preparation practices [(22)] and included in all analyses. Proposed risk factors for food contamination by enteric pathogens included general household conditions that could lead to the introduction (e.g. animals near food) or sustained presence (e.g. floor type) of feces with pathogens in the food preparation and feeding area (Table 1). Due to infrequent (<5%) detection rates for tea, water, and other foods, these types were combined into one “Other” group for the single and multivariable analysis to ensure model convergence.

Analyses were completed through SAS software version 9.4 (SAS Institute, Cary, NC). Separate Generalized linear mixed models (GLMM) were developed for the binary indicator of any pathogen detection (any-path model) and the pathogen diversity measures (sum-path models) to assess relationships between environmental and behavioral risk factors and primary outcomes. For the any-path model, the log link and binomial distribution specifications were used, and regression results were converted to risk ratios. For the Sum-path models, Poisson, zero-inflated Poisson, negative binomial, and zero-inflated negative binomial distributions were evaluated, and the log link and negative binomial distribution family was ultimately determined to best fit the distribution of outcome data. Regression results were converted to risk ratios.

Both sets of models followed the same two-stage process. First, bivariate associations between environmental and behavioral risk factors were determined. Risk factors with p-values smaller than 0.30 in the bivariate testing were included in the multivariable analysis. We then followed a backwards selection process. The Akaike information criterion (AIC) score was noted for the model with all selected-in variables. Then variables were removed individually, and the AIC score was recorded. Variables retained in the final models were selected based upon the model with the lowest AIC score, adjusted for educational level of the caregiver and month of sampling.

## Acknowledgement

This study is collaboration between University of Iowa, Great Lakes University of Kisumu, and the London School of Hygiene & Tropical Medicine. The authors would like to acknowledge University of Iowa’s Environmental Health Sciences Research Center (EHSRC) and the Sanitation and Hygiene for Applied Research Equity (SHARE) consortium for providing the funding. The authors would also like to acknowledge GLUK for helping us in the field implantation. Last, many thanks to the caregivers in the communities of Kisumu, Kenya for participating the study

## Author Contributions

K.K.B, O.C, R.D, J.M and S.S were responsible for study concepts. K.K.B and J.M were responsible for study design. J.M, S.S, and K.T were responsible for carrying out the study. K.K.B and K.T analyzed the data. K.K.B and K.T wrote the manuscript. All authors are responsible for final manuscript editing.

## Supplementary Materials

Table S1: Taqman Array Card Primer and Probes.

Table S2: MS2 Ct Values Across Food Type.

